# Inheritance of neural substrates for motivation and pleasure experience

**DOI:** 10.1101/294090

**Authors:** Zhi Li, Yi Wang, Chao Yan, Ke Li, Ya-wei Zeng, Eric F. C. Cheung, Anna R. Docherty, Pak C. Sham, Raquel E. Gur, Ruben C. Gur, Raymond C. K. Chan

## Abstract

Despite advances in the understanding of the reward system and the role of dopamine in recent decades, the heredity of the underlying neural mechanisms is not known. In the present study, a Monetary Incentive Delay (MID) task was used to examine the haemodynamic activation of the nucleus accumbens (*NAcc*), a key hub of the reward system, in 86 healthy monozygotic twins and 88 dizygotic twins during the anticipation of monetary incentives. The participants also completed self-report measures of pleasure experience. Using a voxel-wise heritability mapping method, activation of the bilateral *NAcc* during the anticipation of monetary gains was found to have significant heritability (*h*^2^ = 0.20-0.49). Moreover, significant shared genetic covariance was observed between pleasure experience and *NAcc* activation when anticipating monetary gain. These findings suggest that *NAcc* activation and self-reported pleasure experience may both be heritable, and their phenotypic correlation may be partially explained by shared genetic variation.

## Introduction

The reward system plays a key role in human behaviour and emotion (*Iversen, 2010*). The nucleus accumbens (*NAcc*), situated in the ventral striatum, is regarded as the hub of the mesolimbic and mesocortical reward systems (*Baldo & Kelley, 2007; Berridge & Robinson, 1998; Haber & Knutson, 2010*). However, the contribution of the *NAcc* to reward processing is not fully understood. Compelling evidence supports the notion that the reward processing system can be parsed into anticipatory and consummatory phases (*Baldo & Kelley, 2007; Craig, 1917*). Dopaminergic activity in the *NAcc* is associated with salience attribution of motivation, which assigns motivational significance to different incentives in the anticipatory period (*Berridge, 2003*, *2007*, *2012*). Despite these advances in understanding the reward system, the heritability of reward processing and its component phenotypes are largely unknown.

Studies in the last decade have suggested a relationship between dopaminergic gene variation and ventral striatal activation measured by functional MRI reward tasks in anticipating, predicting, or receiving monetary incentives (*Aarts et al., 2010; Dreher, Kohn, Kolachana, Weinberger, & Berman, 2009; Forbes et al., 2009; Hahn et al., 2011; Nikolova, Ferrell, Manuck, & Hariri, 2011; Yacubian et al., 2007*). Although single genetic polymorphisms have been associated with ventral striatal activation to a modest extent, studies that examine single or few polymorphisms, are seldom replicated and tend to be underpowered. Evidence from the Psychiatric Genomics Consortium indicates that psychiatric phenotypes are polygenic, with any one gene only contributing a small amount to the overall variance. Thus, to have meaningful explanatory power, it is essential to either use genome-wide common variant data, aggregating the effects of all polymorphisms, or to examine family-based genetic variation. The prediction of ventral striatal activation using polygenic risk scores for psychosis has already revealed the cumulative effect of genes (*Bossong & Kahn, 2016; Lancaster et al., 2016*), and it is expected that related component phenotypes are similarly polygenic. However, the extent to which genetic factors influence *NAcc* and ventral striatal brain activation, particularly in the anticipatory period for reward, is not clearly understood.

In this study, we examined measures of shared genetic variance among dimensional psychiatric phenotypes, using a voxel-wise topographical approach to map striatal activation. While previous studies have mainly employed regions of interest methodology, voxel-wise analysis affords much finer heritability mapping, thereby reducing statistical noise in heritability estimation. Hypoactivation of the *NAcc* and the ventral striatum in anticipating monetary rewards had been reported both in patients with schizophrenia (*Radua et al., 2015*) and in their unaffected biological relatives. Probands and biological relatives also exhibit increased anhedonia that co-segregates in families (*Docherty, Sponheim, & Kerns, 2015; Kendler, Thacker, & Walsh, 1996; Li et al., 2015*). These results suggest that impaired motivation is associated with genetic factors underlying schizophrenia (*de Leeuw, Kahn, & Vink, 2015; Grimm et al., 2014; Vink et al., 2016*). Thus, mapping the heritability of brain activation in anticipating and approaching reward may help clarify its potential role as an endophenotype for psychosis (*Braff, Freedman, Schork, & Gottesman, 2007; Gottesman & Gould, 2003*).

Another important matter to consider is the similar ventral striatal dysfunction reported in patients with major depressive disorder, addiction, and their family members (*Beck et al., 2009; Knutson, Bhanji, Cooney, Atlas, & Gotlib, 2008; Pizzagalli et al., 2009; Wrase et al., 2007*). Since anhedonia and motivational deficits are also relevant to major depressive disorder, the problem of non-specificity needs to be addressed. This study aims to examine the circuitry and behavioral manifestations of dimensional clinical traits, in line with Research Domain Criteria (RDoC) (*Cuthbert, 2014; Cuthbert & Insel, 2010*).

Anhedonia, the diminished ability to experience pleasure, has long been regarded as a core symptom of schizophrenia and major depression disorder (*Meehl, 1975; Meehl, 1990; Pizzagalli, 2014*). Cumulative evidence has also demonstrated a significant heritability estimate for anhedonia ranging from 0.3 to 0.7 (*Kendler & Hewitt, 1992; Linney et al., 2003; MacDonald, Pogue-Geile, Debski, & Manuck, 2001*). Although previous findings support an association between motivation-related *NAcc* activation and anhedonia traits (*Vignapiano et al., 2016; Wacker, Dillon, & Pizzagalli, 2009*), their shared genetic covariance has not been examined. investigating the heritability of *NAcc* activation and anhedonia could facilitate the identification of shared regions of the genome and the exploration of intervention for amotivation and anhedonia, both of which indicate poor prognosis and are refractory to currently available treatment (*Kring & Barch, 2014*).

In the present study, we adopted a healthy-twin design to examine the heritability of pleasure experience and its shared genetic covariance with *NAcc* activation during the anticipation of monetary rewards using the Monetary Incentive Delay Task (MID). We hypothesized that both motivation-related *NAcc* activation and pleasure experience would exhibit significant heritability. We further hypothesized that motivation-related *NAcc* activation and pleasure experience would share significant genetic covariance.

## Results

### Demographics

The sample included 43 pairs of same-sex monozygotic (MZ) twins and 44 pairs of same-sex dizygotic (DZ) twins, who were matched in gender ratio, age, and years of education. In addition, their head motion parameters were also comparable (Table 1). The MZ and DZ twins demonstrated comparable scores on the Temporal Experience Pleasure Scale (TEPS), whereas DZ twins scored lower than MZ twins on the revised Chinese versions of Chapman Social Anhedonia Scale (RCSAS) and Chapman Physical Anhedonia Scale (RCPAS). The ICC of RCPAS and TEPS scores among the MZ twins were twice that of the DZ twins. However, the ICC of RCSAS scores among MZ and DZ twins did not significantly vary from 0 (Table 1). These findings suggest that in this healthy sample, there was insufficient power to detect significant genetic or phenotypic variation in RCSAS scores.

**Table 1.** Demographics of participants

| | MZ (N = 86) | ICC | DZ (N = 88) | ICC | $\chi^2/t/F$ | p | ALL (N = 174) |
| --- | --- | --- | --- | --- | --- | --- | --- |
| Gender (Male/Female) | 46/40 | / | 46/42 | / | 0.026 | 0.872 | 92/82 |
| Age | 19.98(1.89) | / | 19.8(1.76) | / | -0.66 | 0.513 | 19.89(1.82) |
| Education | 12.33(1.75) | 0.671** | 12.44(1.63) | 0.591** | 0.46 | 0.647 | 12.39(1.69) |
| FD | 0.13(0.04) | 0.204 | 0.13(0.04) | -0.059 | 0.156 | 0.694 | 0.13(0.04) |
| RCSAS | 11.38(5.96) | 0.087 | 7.57(4.48) | 0.281 | 22.38 | < 0.001** | 9.45(5.59) |
| RCPAS | 22.12(8.03) | 0.490** | 18.97(7.66) | 0.201 | 6.29 | 0.013* | 20.52(7.98) |
| TEPS | 4.06(0.59) | 0.352* | 4.08(0.59) | -0.02 | 0.004 | 0.947 | 4.07(0.59) |
| TEPS_Anticipation | 4.11(0.67) | 0.396** | 4.13(0.7) | -0.021 | 0.19 | 0.662 | 4.12(0.68) |
| TEPS_Consummation | 4.11(0.77) | 0.346* | 4.12(0.75) | 0 | 0.19 | 0.667 | 4.14(0.76) |

Note: \*, p < 0.01; \*\*, p < 0.05; MZ = Monozygotic twins; DZ = Dizygotic twins; ICC = Interclass correlation coefficient; FD = Frame Displacement; RCSAS = Revised Chapman Social Anhedonia Scale; RCPAS = Revised Chapman Physical Anhedonia Scale; TEPS = Temporal Experience Pleasure Scale.

### Brain activation in contrast [Gain vs. No-incentive] of MID task

For this contrast, during the anticipatory phase, there was significant activation of the bilateral *NAcc* and the thalamus in all participants. Activation of the bilateral *NAcc* was also observed in MZ and DZ twins respectively. Furthermore, activation of the left insula was observed in MZ twins, whereas activation of the right insula was observed in DZ twins (Table 2 & Figure 1B).

**Table 2.**
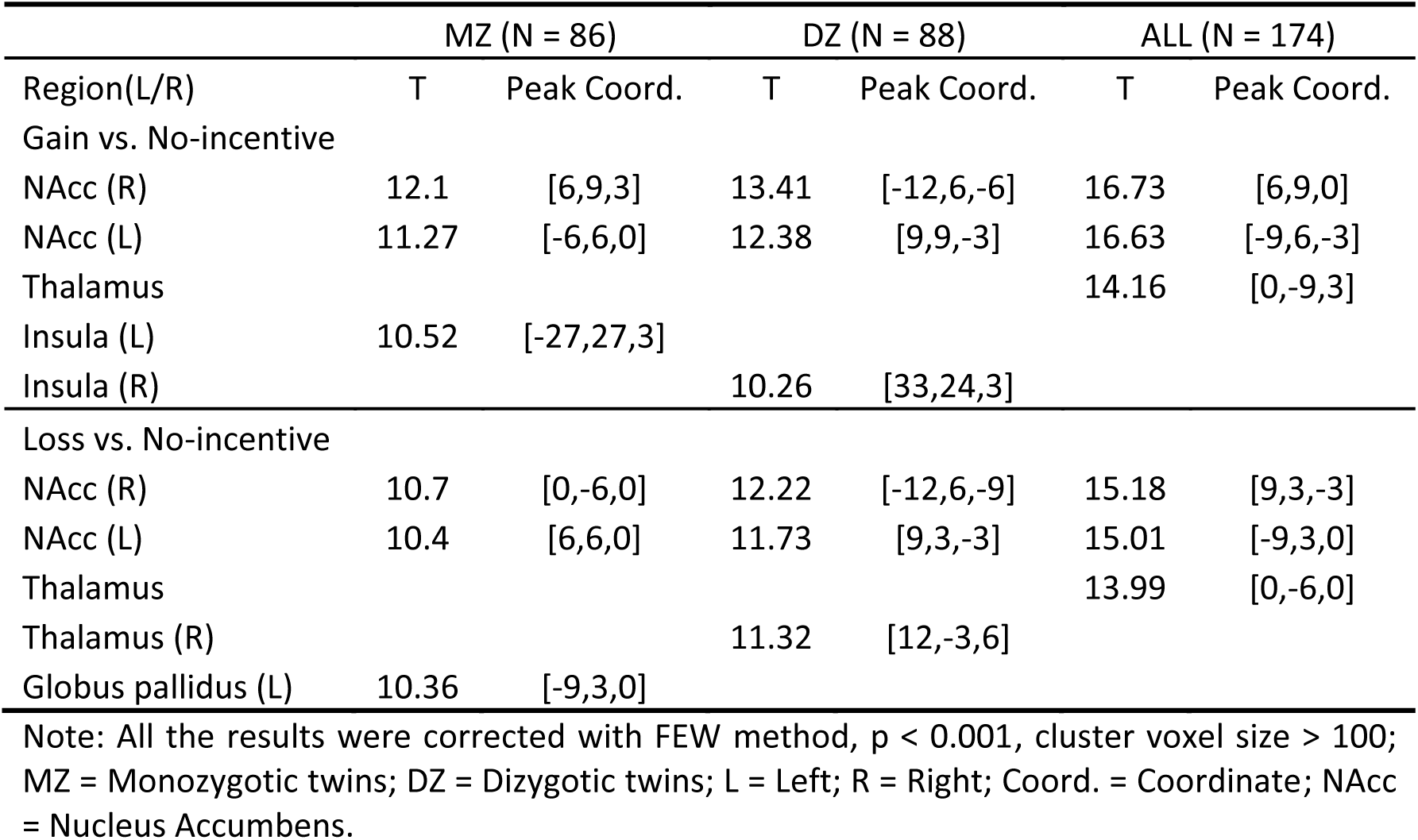
Whole brain activation in anticipating monetary incentive.

**Figure1.**
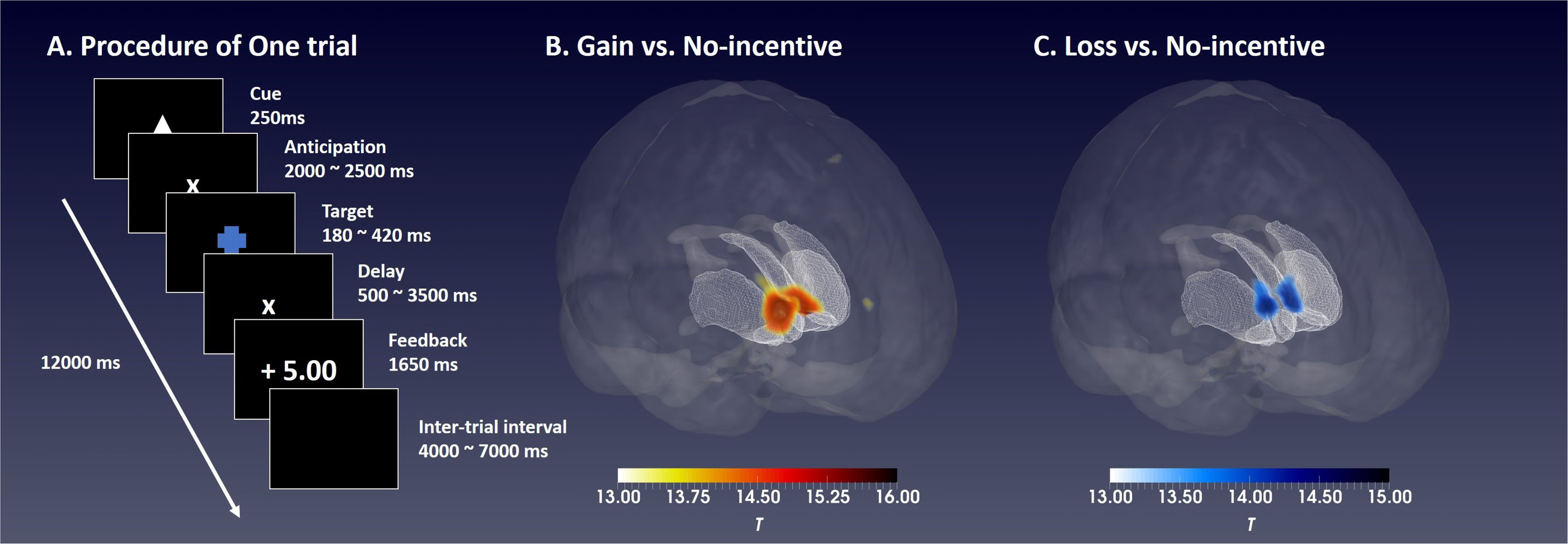
Monetary incentive delay (MID) task and brain activation in anticipating monetary gain and loss. A: The Workflow of one trial in the MID task; B: The activation of bilateral nucleus accumbens (*NAcc*) was found in contrast [Gain vs. no-incentive] during the anticipatory phase, the white-red color bar indicates the statistical t value; C: The activation of bilateral *NAcc* was found in contrast [Loss vs. no-incentive] during the anticipatory phase, the white-blue color bar indicates the statistical t value.

### Brain activation in contrast [Loss vs. No-incentive] of MID task

Significant activation of the bilateral *NAcc* and the thalamus were observed in the contrast [Loss vs. No-incentive] during the anticipatory phase in all participants. Significant activation of the bilateral *NAcc* was observed in both MZ and DZ twins, while activation of the left globus pallidus and the right thalamus was observed in MZ twins and DZ twins respectively (Table 2 & Figure 1B).

### Heritability brain mapping

In the voxel-by-voxel heritability brain mapping, two clusters were detected with significant heritability in the contrast [Gain vs. No-incentive]. The right one contained 33 voxels, while the left one contained 15 voxels. The coordinates of the peak point of the two clusters were [9,15,-3] and [−9,18,-6], which were located at the right *NAcc* and the left *NAcc* respectively (Figure 3). The *h*^2^ of each voxel within both clusters ranged from 0.2 to 0.49, and the average *h*^2^ was 0.34 (Figure 3). However, we did not find any significant heritability for the contrast [Loss vs. No-incentive].

### Genetic model fitting

As shown in Table 3, the Cholesky trivariate model (AIC = 382.4) was not worse than the saturated model (AIC = 406.45) (Δ− 2LL = 41.94, Δ df = 33, *p* = 0.14). The nested sub-models were compared with the trivariate model. A marginally significant deterioration emerged when all the additive genetic paths (a11, a12, a13, a22, a23, a33) were deleted (Δ− 2LL = 12.59, Δ df = 6, *p* = 0.05), whereas the model did not worsen when we discarded all the common environmental paths (c11, c12, c13, c22, c23, c33) (Δ− 2LL = 0.25, Δ df = 6, *p* = 1). Results of model comparison indicated that additive genetic factors, rather than common environmental factors, significantly contributed to the full model. The trivariate AE model (AIC = 370.65) was adopted for the subsequent parameter dropping and model comparison. After dropping one path, Model X (with path e21 dropped) was found to have the smallest AIC (368.66) in its class and was hence adopted (Δ− 2LL = 0.01, Δ df = 1, *p* = 0.94). Based on Model X, a second path was further discarded. Finally, Model XV (with paths e21 and e31 dropped) had the smallest AIC (367.35) among all the tested models and was selected as the best model (Δ− 2LL = 0.69, Δ df = 1, *p* = 0.4). A third path was further dropped based on Model XV, but all the sub-models significantly deteriorated compared with Model XV.

**Table 3.** Model fitting and selection

| No. | Model | ep | -2LL | df | AIC | diffLL | diffdf | *p* |
| --- | --- | --- | --- | --- | --- | --- | --- | --- |
| I | Saturated Model | 54 | 1342.45 | 468 | 406.45 | / | / | / |
|  | **Cholesky Trivariate Model** | **21** | **1384.4** | **501** | **382.4** | **41.94** | **33** | **0.14** |
| II | Drop a or c matrix |  |  |  |  |  |  |  |
| III | **Drop c matrix** | **15** | **1384.65** | **507** | **370.65** | **0.25** | **6** | **1** |
| IV | Drop a matrix | 15 | 1396.98 | 507 | 382.98 | 12.59 | 6 | 0.05 |
|  | Drop one path based on the model III |  |  |  |  |  |  |  |
| V | Drop a31 | 14 | 1387.21 | 508 | 371.21 | 2.56 | 1 | 0.11 |
| VI | Drop a32 | 14 | 1387.68 | 508 | 371.68 | 3.03 | 1 | 0.08 |
| VII | Drop a21 | 14 | 1390.61 | 508 | 374.61 | 5.95 | 1 | 0.01 |
| VIII | Drop e32 | 14 | 1390.07 | 508 | 374.07 | 5.42 | 1 | 0.02 |
| IX | Drop e31 | 14 | 1385.22 | 508 | 369.22 | 0.57 | 1 | 0.45 |
| X | **Drop e21** | **14** | **1384.66** | 508 | 368.66 | 0.01 | 1 | 0.94 |
|  | Drop two paths based on the model X |  |  |  |  |  |  |  |
| XI | Drop e21 and a31 | 13 | 1387.37 | 509 | 369.37 | 2.71 | 1 | 0.1 |
| XII | Drop e21 and a32 | 13 | 1388.4 | 509 | 370.4 | 3.75 | 1 | 0.05 |
| XIII | Drop e21 and a21 | 13 | 1393.18 | 509 | 375.18 | 8.53 | 1 | 0 |
| XIV | Drop e21 and e32 | 13 | 1390.07 | 509 | 372.07 | 5.42 | 1 | 0.02 |
| XV | **Drop e21 and e31** | **13** | **1385.35** | **509** | **367.35** | **0.69** | **1** | **0.4** |
|  | Drop three paths based on the model XV |  |  |  |  |  |  |  |
| XVI | Drop e21 and e31 and a31 | 12 | 1393.29 | 510 | 373.29 | 7.94 | 1 | 0 |
| XVII | Drop e21 and e31 and a21 | 12 | 1394.03 | 510 | 374.03 | 8.68 | 1 | 0 |
| XVIII | Drop e21 and e31 and a32 | 12 | 1388.44 | 510 | 368.44 | 3.09 | 1 | 0.08 |
| XIX | Drop e21 and e31 and e32 | 12 | 1390.56 | 510 | 370.56 | 5.21 | 1 | 0.02 |

Note: No. = Model number; Model = Parameters drop by this model; ep = estimate parameter; -2LL = minus 2 log likelihood; df = degree of freedom; AIC = Akaike information criterion; diffLL = the difference of minus 2 log likelihood between two models; diffdf = the difference of the degree of freedom between two models; *p* indicates the significance of model deterioration between two models. The trivariate genetic model was compared with the saturated model first, then the sub-models II – IV were compared with the full genetic model from which the model III was selected as the best-fitted model for the next step parameter dropping. One parameter was first drop from the model III in which the model X was selected. Then two parameters were drop based on the model X. Finally, the model XV with the smallest AIC among all tested sub-models was chosen as the best fitted model, which model did not deteriorate by compared with the model X (*p* > 0.05).

In the best-fitting Model XV (Figure 4), the *h*^2^ of bilateral *NAcc* activation during the anticipation of monetary gain was 0.43 [95% Confidential Interval (CI): 0.19-0.62], the *h*^2^ of RCPAS scores was 0.61 (95%CI: 0.39-0.75), and the *h*^2^ of TEPS scores was 0.29 (95%CI: 0.01-0.54). *NAcc* activation shared the same genes with RCPAS (*r*_*g*_ = - 0.45) and TEPS (*r*_*g*_ = 0.59) scores. In addition, scores on the RCPAS and the TEPS were also influenced by some of the same genes (*r*_*g*_ = −0.72). Addictive genetic factors contributed almost 100% to the phenotypic correlation between *NAcc* activation and RCPAS scores (*r*_*ph*_ = −0.23, *r*_*ph(g)*_ = −0.23) and between *NAcc* activation and TEPS scores (*r*_*ph*_ = 0.21, *r*_*ph(g)*_ = 0.21). Sixty-four percent of the phenotypic correlation between RCPAS and TEPS scores (*r*_*ph*_ = −0.47) was attributed to additive genetic factors (*r*_*ph(g)*_ = −0.3), while 36% of the correlation was attributed to environmental factors (*r*_*ph(e)*_ = −0.17).

## Discussion

This is the first biometrical study to examine the heritability of neural substrates underlying reward processing and the shared genetic covariance of motivation-related *NAcc* activation and pleasure experience. Consistent with our hypothesis, activation in the bilateral *NAcc* during the anticipation of monetary gain was significantly heritable. The heritability estimate of each voxel within the bilateral *NAcc*, while low-to moderate, was significant and ranged from 0.2 to 0.49, whilst the heritability estimate of *NAcc* activation as a whole was 0.43. RCPAS and the TEPS also exhibited significant heritability in healthy twins, at 0.61 and 0.29 respectively. Even with a modest sample size for ACE modeling, motivation-related *NAcc* activation evidenced significant shared genetic covariation with both self-reported measures of pleasure experience. This suggests that the significant phenotypic correlation between *NAcc* activation and pleasure experience is partially accounted for by shared genetic variation.

Bilateral *NAcc* activation during the anticipatory phase for monetary incentives, found in our whole brain analysis, is consistent with previous findings (*Knutson, Fong, Adams, Varner, & Hommer, 2001; Knutson & Greer, 2008; Knutson, Westdorp, Kaiser, & Hommer, 2000*), which supports the validity of the MID task in correlating with *NAcc* activation *in vivo*. Evidence from animal studies supports the role of dopamine within the *NAcc* in salience attribution, an essential component of motivation and goal-directed behaviour formulation (*Berridge, 2007*, *2012; Berridge & Robinson, 1998*). Although fMRI measures brain haemodynamics rather than neurotransmitter chemistry, previous studies have linked activation at the *NAcc* and the ventral striatum in rewarding tasks to local dopaminergic activity. Amphetamine-induced dopamine release at the *NAcc* has been associated with local haemodynamic activation in reward processing (*Knutson & Gibbs, 2007; Oswald et al., 2007*). These data suggest that *NAcc* activation measured through fMRI could indirectly reflect local dopaminergic activity. In addition, polymorphisms of dopamine have been associated with ventral striatal activation in reward processing, suggesting that facets of the mesolimbic reward system may be heritable (*Dreher et al., 2009; Forbes et al., 2009; Yacubian et al., 2007*). Indeed, significant MZ twin correlation in *NAcc* activation has previously been reported (*Silverman et al., 2014*). In the present study, we corroborated previous findings through quantifying the genetic and environmental effects on the motivation-related *NAcc* activation. Stokes and colleagues (2013) reported a significant heritability estimate of 0.21 for dopaminergic activities of the right ventral striatum in the resting state. In this study, we found a heritability estimate of 0.4 for bilateral *NAcc* activation in anticipating monetary gains. The considerably larger sample size in this study, relative to previous research, lends further support to this finding.

Quantifying the heritability of *NAcc* activation in anticipating monetary incentives could facilitate our understanding of the genetic effect on reward processing and detection of associated genetic loci. Voxel-wise heritability estimation is an alternative methodology that may be more sensitive in detecting genetic effects, and as such may supplement findings from previous regions of interest analysis often limited by small sample sizes. Across single-gene polymorphism studies (*Dreher et al., 2009; Forbes et al., 2009; Yacubian et al., 2007*), twin studies (*Silverman et al., 2014*) or studies estimating the heritability of striatal dopaminergic activity (*Stokes et al., 2013*), all of which adopted the mean value within a region of interest (ROI), a lateralization bias toward the right *NAcc* or ventral striatal activation has been reported. However, such lateralization pattern was not observed in the present voxel-wise analysis. This is consistent with a voxel-wise heritability approach being more sensitive in detecting the heredity of brain activation compared with traditional ROI analysis.

It should be noted that only the *NAcc* activation in anticipating monetary gain, rather than loss, showed significant heritability. This result is consistent with previous findings reporting significant correlation between *NAcc* activation in older and younger MZ twins in anticipating monetary gain but not loss (*Silverman et al., 2014*). One possible explanation is that there exists less individual difference in anticipating monetary gain than loss, even though the *NAcc* activation appears to be sensitive to both incentive conditions. The small or non-significant heritability could be attributed to the high phenotypic variance within both MZ and DZ twins due to limited sample size. It is notable that the studies mentioned above only linked dopaminergic gene polymorphism to ventral striatal activation in processing monetary gain but not loss, which deserves further investigation.

We also investigated the heritability of pleasure experience. In this study, physical anhedonia traits measured by the RPCAS also demonstrated significant heritability, whilst experiential pleasure measured by the TEPS was characterized by moderate but significant heritability. These findings are consistent with previous studies (*Hay et al., 2001; Kendler & Hewitt, 1992; Linney et al., 2003*). In these studies, however, significant heritability has also been detected in social anhedonia which was not found in our study. Cultural factors may be a possible influence accounting for this difference, but this requires further investigation.

In the best-fitting genetic model, *NAcc* activation during the anticipation of monetary gain exhibited significant genetic share with scores on the RPCAS and the TEPS. In addition, the phenotypic correlation between *NAcc* activation and pleasure experience was almost entirely attributed to genetic factors. These data suggest that motivation-related *NAcc* activation during the anticipation of monetary rewards may share common additive genes with pleasure experience, and these genes contribute significantly to the phenotypic expression. Locating the shared genes of these two phenotypes could facilitate clarification of the underlying neurobiological mechanisms of amotivation and anhedonia in schizophrenia and major depressive disorder.

The main limitation of this study is the modest sample size, which was relatively small for statistical methods applying ACE models. While 196 participants with 174 valid data sets could be regarded as a medium to large sample in task-based fMRI research, these preliminary results must be replicated in a larger cohort. Importantly, adopting voxel-wise analysis with correction for multiple comparisons could partially compensate for the relatively small sample size. One primary problem in genetic modeling lies in within-group variation, which is sensitive to the presence of outliers; however, in this study we detected none. Furthermore, demographic variables and head movements were carefully matched among subgroups with an aim to enhance the validity of our findings.

In conclusion, our findings suggest that motivation-related *NAcc* activation in anticipating monetary gain and pleasure experience are at least partially heritable. Importantly, motivation-related *NAcc* activation and pleasure experience exhibit significant shared genetic covariance. Future molecular studies examining shared polygenicity of these traits would further inform this research. Locating and refining areas of the genome associated with expression of these traits will ultimately aid in the understanding of the underlying neurobiological mechanisms of treatment-refractory symptoms.

## Materials and methods

### Participants

Forty-seven pairs of same-sex dizygotic (DZ) twins and 51 pairs of same-sex monozygotic (MZ) twins were recruited from the Twin Registry of the Institute of Psychology, the Chinese Academy Sciences (*Chen et al., 2010; Chen et al., 2013*). The zygosity of each twin was jointly determined by DNA analysis based on saliva, and two zygosity questionnaires (*Chen et al., 2010*). Potential participants were excluded from the study if they: 1) had a personal or family history of diagnosable mental disorders; b) had a history of head trauma or encephalopathy; 3) had a history of substance abuse, including tobacco and alcohol; 4) had an Intelligence Quotient (IQ) lower than 70; 5) had severe hearing or visual impairment; or 6) were ambidextrous or left-handed. This information was verified by the Twin Registry, the participants themselves, and their guardians. Detailed experimental procedures conformed to the Declaration of Helsinki and all participants gave written informed consent. All participants completed checklists capturing experiential pleasure and hedonic traits before the brain scans. They then undertook the Monetary Incentive Delay (MID) task inside the scanner. Each participant received 450 RMB as compensation. The study was approved by the Ethics Committee of the Institute of Psychology, the Chinese Academy of Sciences.

### Self-report measures of pleasure experience

The revised Chinese versions of the Chapman Physical Anhedonia Scale (RCPAS) and the Chapman Social Anhedonia Scale (RCSAS) were administered to measure physical and social anhedonic traits, respectively (*Chan, Wang, et al., 2012*). The RCPAS consists of 61 true-false items, whereas the RCSAS consists of 40 true-false items. These two scales have been stable and valid in measuring anhedonic traits across time and along the schizophrenia spectrum (*Chan, Gooding, et al., 2016*). The Chinese version of the Temporal Experience Pleasure Scale (TEPS) is a 19-item checklist that was used to measure anticipatory and consummatory pleasure in each participant (*Chan, Shi, et al., 2012*). The Chinese version has good psychometric properties for measuring anhedonia across the different stages of schizophrenia (*Chan et al., 2010; Chan, Shi, et al., 2012; Li et al., 2015*).

### Monetary Incentive Delay (MID) task

We adopted the original version of the MID task developed by Knuston and colleagues (*Knutson et al., 2000*) to the abridged imaging version (*Chan, Li, et al., 2016*). In each trial of the task, a 250-millisecond cue indicating three different conditions was first presented at the centre of the screen, followed by a blank interval that jittered between 2000 and 2500 milliseconds. Then a blue target with adjusted duration was displayed, followed by an interval that jittered between 500 and 3500 milliseconds. Finally, a 1650-millisecond feedback was presented followed by an inter-trial interval that jittered between 4000 and 7000 milliseconds. Each trial lasted 12000 milliseconds (Figure 1A).

Participants were asked to hit the target as quickly as possible by pressing the right button on a panel with their right thumb. The initial duration of the target was set at 300 milliseconds and changed according to the subsequent performance of each participant. If a target was successfully hit twice, the target duration was reduced by 20 milliseconds. Alternatively, if a target was missed twice, 20 milliseconds were added to the target duration. Using this strategy, the hit rate of each participant was controlled at around 66.7%. The three cues indicated three different conditions: a triangle indicated a monetary gain condition, a square indicated a monetary loss condition, and a circle indicated a no-incentive condition. Participants gained five monetary points if they hit the target in the monetary gain condition, or lost five monetary points if they missed the target in the monetary loss condition. In the no-incentive condition, participants did not gain or lose any points whether the target was hit or not. Two runs containing 10 gain, 10 loss and 10 no-incentive conditions in each run were applied. The trials in each run were presented in a pseudo-random order. Participants practiced with an independent 30-trial run before entering the scanner and were informed that their final monetary points gained in the scanner could be converted into cash and added to their compensation. This abridged version has been shown to activate the *NAcc* effectively in healthy, subclinical and clinical samples (*Chan, Li, et al., 2016; Smoski, Rittenberg, & Dichter, 2011*).

### Brain image acquisition

Brain imaging data were collected with 32-channel head coil in a 3 T Siemens Trio MRI Scanner. An experienced radiologist who was blind to this study was responsible for data acquisition. A T2-weighted FLAIRE sequence (TR = 4000ms; TE = 90ms; FOV = 200mm^2^; slices = 19; flip angle = 120°; image matrix = 256×512; voxel dimensions = 0.9 × 0.4×5mm^3^) was first used to ascertain that each participant had no organic brain disorders. Then a gradient-echo echo-planner sequence (TR = 2000ms; TE = 30ms; FOV = 210mm^2^; slices = 32; flip angle = 90°; image matrix = 64×64; voxel dimensions = 3.3 × 3.3 × 4mm^3^; Number of TR = 184 for each run) was applied to acquire the functional brain activation imaging data of each participant while performing the MID task. Finally, a high resolution structural brain image was acquired for anatomical registration (TR = 2300ms; TE = 3ms; FOV = 256mm^2^; flip angle = 9°; image matrix = 256×256; voxel dimensions = 1 × 1 x 1mm^3^). Each participant wore earplugs during scanning. Their heads were fixed with a vacuum pillow and sponge pads to minimize head motion.

### Imaging data processing

The latest version of the SPM 12 (Wellcome Trust Centre for Neuroimaging) was used for imaging data processing. The functional images were first realigned into the first volume of each scanning sequence for movement correction, followed by slice timing correction. The observed head motion parameters, three transitions and three rotations, were calculated into the framewise displacement (FD), a comprehensive and reliable index for head movement (*Power, Barnes, Snyder, Schlaggar, & Petersen, 2012*). Participants with maximum head motion higher than 2 mm and 2 degrees, and mean FD larger than 0.25 mm were excluded from the final analysis, along with their twin. Individual high-resolution brain structure image was non-linearly registered to the Montreal Neurological Institute (MNI) template that produced a transformation matrix. Using this matrix, all functional brain images were normalized into a common standard atlas. Functional images were resampled into 3 × 3 x 3 mm^3^ and spatially smoothed with a 6mm full-width-at-half-maximum (FWHM) Gaussian isotropic kernel. A 128 Hz high-pass filter was applied to the time series of each voxel to remove low-frequency noise.

The preprocessed functional imaging data were included in a first-level general linear model (GLM) with three predictors of interest during the anticipatory phase for monetary incentives: gain, loss, and no-incentive. First, the data for each participant was analyzed to provide a voxel-wise t-statistic map for each contrast, [gain vs. no-incentive] and [loss vs. no-incentive], during the anticipatory phase. The time points of target hitting and feedback period were included as parametric modulation to minimize their influence on the anticipation for incentives. Six raw head movement parameters were also included as covariates to remove motion effect. For each contrast, the t-statistics map of all the participants were included into a GLM with the t-statistics as the dependent variable and the FD as covariate to further minimize head motion effect. To clarify whether the bilateral *NAcc* were activated in both MZ and DZ twins, one-sample t-tests were also applied to the t-statistics of the MZ and DZ groups. The statistical significance threshold of the whole brain analysis was set as p < 0.001 with FWE correction and cluster voxel size > 100. Since 11 pairs of twins were excluded due to excessive head movements, 44 pairs of DZ twins and 43 pairs of MZ twins were included in the final analysis.

### Statistical analysis

Pearson chi-square test was used to test if the gender ratio of MZ and DZ twins were different from each other. The matching of age and years of education between MZ and DZ twins was tested with independent t-test. Univariate general linear model with gender, age, and years of education as covariates was used to compare FD, scores on the RCSAS, the RCPAS, the TEPS and their various subscales between MZ and DZ twins. The intraclass correlation coefficients (ICC) of behavioral phenotypes among the MZ and DZ twins were calculated respectively. If the ICC of a phenotype among MZ twins was significantly twice larger than that for DZ twins, the phenotype in question may be influenced by familial factors.

### Heritability brain mapping

Voxel-by-voxel heritability brain mapping was carried out with the latest version of the OpenMx (*Neale et al., 2016*), FSL (*Jenkinson, Beckmann, Behrens, Woolrich, & Smith, 2012*) and in-house MATLAB (The MathWorks, Inc., Natick, Massachusetts, United States) scripts. Since mapping the heritability of all voxels in the whole brain may increase the possibility of type II error, we adopted the Oxford-GSK-Imanova structural striatal atlas which contained the core brain regions sensitive to dopaminergic activity and reward tasks (*Tziortzi et al., 2011*). The structural striatal template was first resampled into a 3 × 3 x 3 mm^3^ mask containing 765 voxels. The T values of voxels in the striatal mask were extracted from the two contrast files, [gain vs. no-incentive] and [loss vs. no-incentive] of each participant. There were no outliers more than three standard deviations from the mean T value. Age, gender, years of education, and FD were regressed out from the extracted t values to remove their possible influences on heritability estimation (*Bergen, Gardner, & Kendler, 2007; Lenroot et al., 2009*). Finally, the standardized residuals were submitted to the genetic model. A conventional univariate ACE model was used, in which A denotes additive genetic effects, C denotes common environmental effects, and E denotes unique environmental effects. MZ twins are assumed to share 100 percent of the additive genetic variance and common environmental variance, whereas the DZ twins are assumed to share 50 percent of the additive genetic variance and 100 percent of common environmental variance. The part accounted for by A in the total variance was defined as the heritability estimate *h*^2^ of this phenotype (*Neale & Maes, 2004*). To clarify the significance of components A, C and E, sub-models AE and CE were compared to the full ACE model, and model E was compared to AE and CE models respectively. If model fit significantly decreased, then the dropped factor was considered essential in the model. In model selection, the model with the smallest Akaike’s Information Criterion (AIC) value was selected as the best-fit model. We adopted the *p* value of model comparison between the AE and the E model to test the significance of *h*^2^ if the AE model was detected as the best-fit model the full ACE model failed to surpass its sub-models in any voxel from the Oxford-GSK-Imanova structural striatal atlas) (*Neale & Maes, 2004*). Finally, FDR correction with adjusted p < 0.05 was applied to the acquired p-maps to adjust for multiple comparisons. Comparing to the full ACE model in brain functional or structural studies, the method in the present study allowed us to quantify the heritability of brain activation and relevant significance in the best-fit genetic model, and to correct for multiple tests (*Li et al., 2018*). The cluster tool of FSL was used to identify clusters in which *h*^2^ was significant. Masks with voxels in which the 95% confidence interval of *h*^2^ did not contain zero were also added onto the brain heritability map.

### Multivariate Model fitting

A trivariate Cholesky ACE model was fitted to examine the genetic sharing between *NAcc* activation and pleasure experience (RCPAS and TEPS scores) (Figure 2). T values of the contrast [gain vs. no-incentive] were extracted from bilateral *NAcc* mask with significant heritability from the heritability brain mapping step. Gender, age, and years of education were then regressed from the extracted mean T value, RCPAS, and TEPS scores. The FD value was additionally regressed from the extracted mean T value for head motion correction. The *NAcc* activation during the anticipation of monetary loss failed to show significant heritability. In addition, MZ twin correlation in social anhedonia was not higher than DZ twin correlation and both correlation coefficients were not significant. Hence, The *NAcc* activation in contrast [loss vs. no-incentive] and RCSAS were not included in the multivariate genetic model presented in Figure 2. The trivariate ACE model was first compared with the saturated model in which all the constraints were set as free. Then the nested sub-models were compared with the full trivariate ACE model in a stepwise manner. We first discarded all the model paths containing A, C or E effect in a stepwise manner (Figure 2 & Table 3). After that we discarded one path from the best fitted model observed from the above step, and then two paths until the model significantly deteriorated (indicated by *p* < 0.05). The model with the smallest AIC was selected as the best-fit model, and the likelihood ratio chi-squared statistic, minus two times log likelihood difference (–2LL) were applied for model comparison. The contribution of A, C, and E to the phenotype correlation between two phenotypes were estimated using the following formulae: 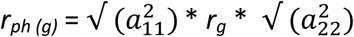;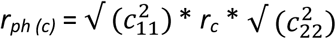;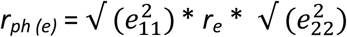 (*Toulopoulou et al., 2015*).

**Figure2.**
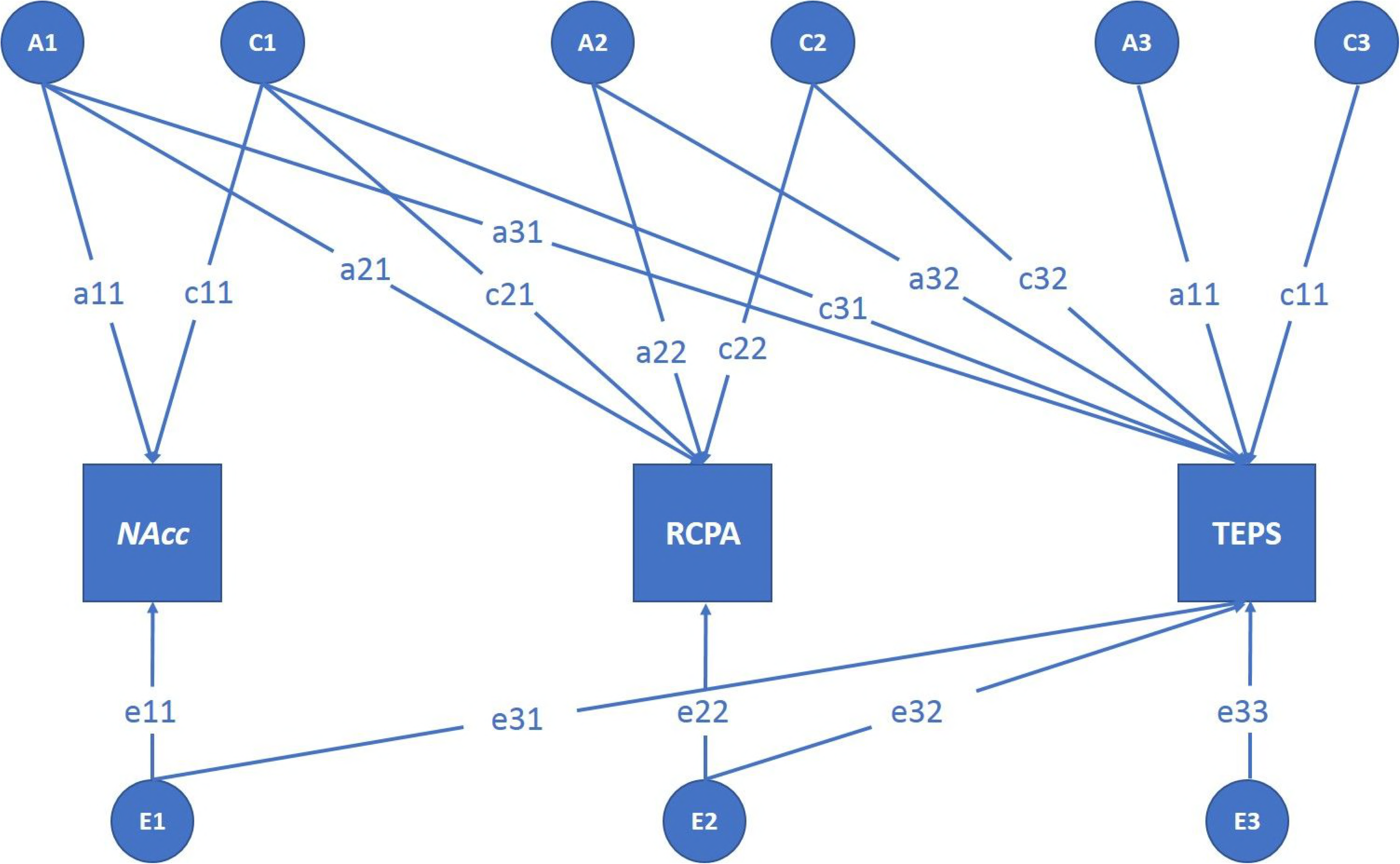
Full trivariate genetic model with Cholesky decomposition. The *NAcc* indicates the extracted t value from the contrast file of [Gain vs. No-incentive], with the mask from the voxel-wise heritability mapping of brain activation. The RPCAS indicates the physical anhedonia trait, whilst the TEPS indicates the experiential pleasure. The A indicates the additive genetic effect, the C indicates the common environmental effect, and the E indicates the unique environmental effect. The lowercase with number indicates the path of model, such as ‘a21’ indicates the influence from the additive gene effect of *NAcc* to RPCAS.

**Figure3.**
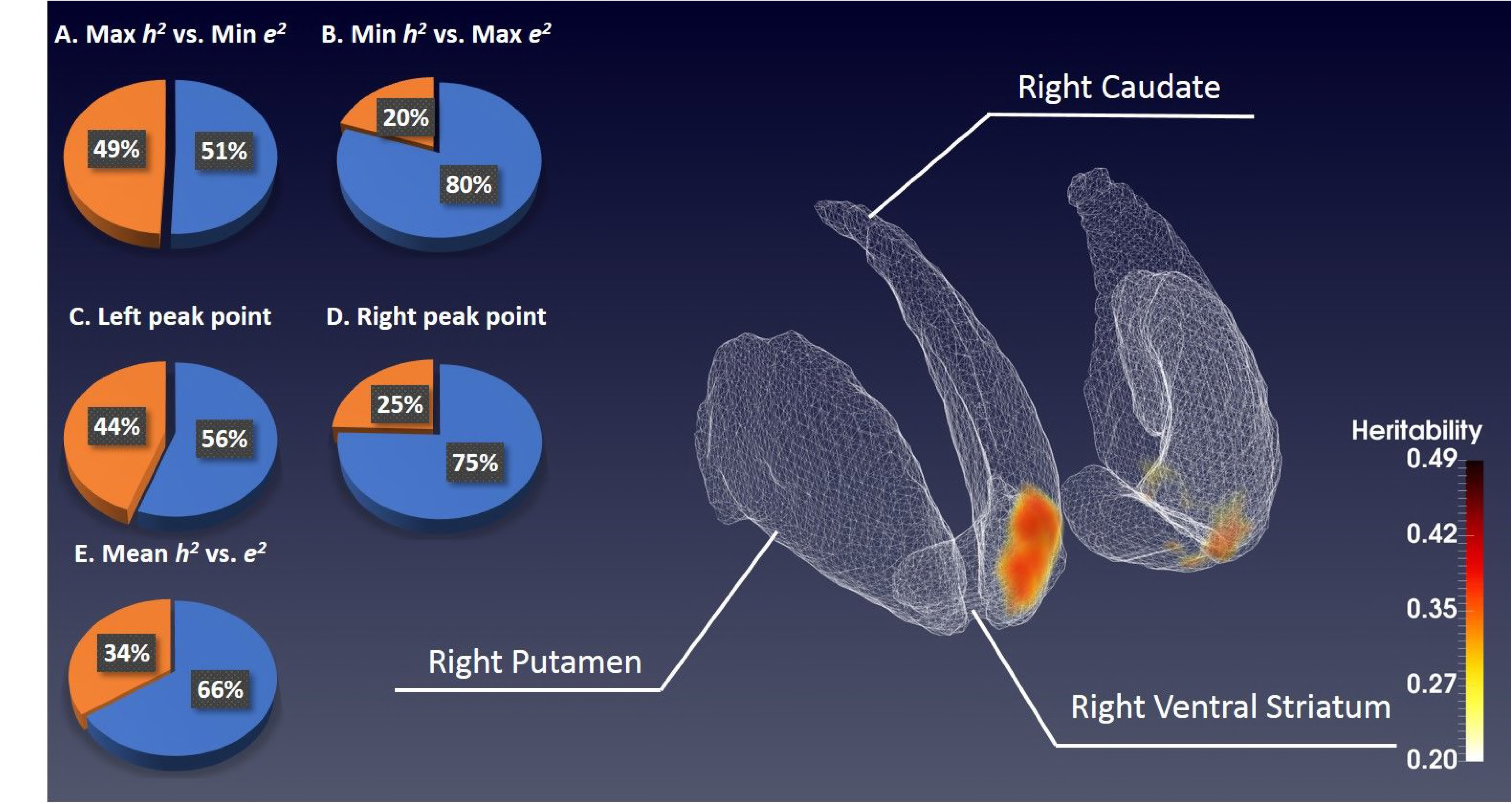
Voxel-wise heritability mapping on brain activation. In the standard striatum atlas on the right, two clusters with significant heritability in bilateral *NAcc* are detected, which ranges from 0.2 to 0.49 in each voxel (reflected by the white-red color bar). The five pie charts on the left indicate the heritability and unique environmental effect of AE model on the two clusters, in which the golden color indicates the heritability whilst the blue color indicates the unique environmental effect.

**Figure4.**
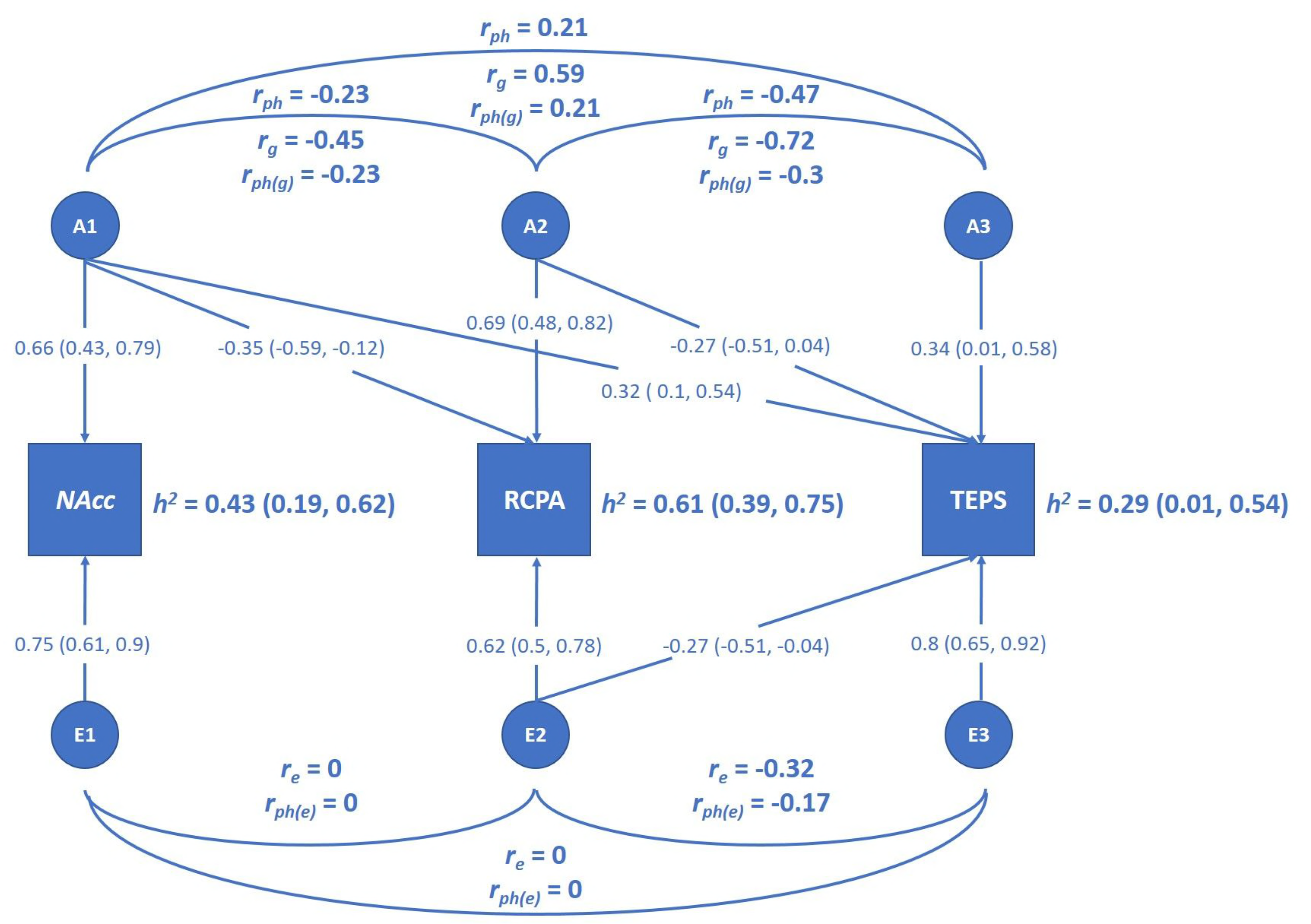
The best-fitted trivariate genetic model. Only the additive genetic and unique environmental effects contribute to this model. The *h*^2^ indicates the heritability with 95% confidence interval of each single phenotype, the *r*_*ph*_ indicates the phenotypic correlation between phenotypes, the *r*_*g*_ indicates the genetic correlation, and the *r*_*e*_ indicates the unique environmental correlation. The *r*_*ph(g)*_ and r_ph(e)_ indicates the contributions to phenotypic correlation, from the additive gene and unique environment respectively. The standard path coefficients with 95% confidence interval were also supplied in this figure.

## Acknowledgements

This study was supported by a grant from the Beijing Municipal Science & Technology Commission Grant (Z161100000216138), National Key Research and Development Programme (2016YFC0906402), the National Science Fund China (81571317), the “Strategic Priority Research Programme (B)” of the Chinese Academy of Sciences (XDB02030002), and the CAS/SAFEA International Partnership Programme for Creative Research Teams (Y2CX131003). The authors would like to thank Ting Xu, Qing, Zhao, Chao Yan, Hui-jie Li, Yu-na Wang, Yan-fang Shi, Xiao-yan Cao, Weizhen Xie, Jie Chen, Xin-ying Li, Jie Zhang and the team members of the Twins Registry of Institute of Psychology, Chinese Academy Sciences for their kind recruitment of healthy twins, and all the twin participants for their taking part in the present study.

## Author contributions

Zhi Li designed the study, collected and analyzed the data, and wrote up the manuscript. Yi Wang, Chao Yan, Ke Li and Ya-wei Zeng collected all the data and commented on the first draft of the manuscript. Eric F. C. Cheung, Anna R. Docherty, Pak C. Sham, Raquel E. Gur and Ruben C. Gur provided significant comments to the manuscript. Raymond C. K. Chan conceptualized the idea, designed the study, interpreted the findings and commented the manuscript critically.

## Additional files

Figure 1 & Table 2 –Source Data 1_[loss vs. no-incentive]

Figure 1 & Table 2 –Source Data 2_[gain vs. no-incentive]

Figure 3-Source Data

Figure 4 & Table 3 -Source Data 1

Figure 4-& Table 3 Source Data 2

Raw data 1_Activation in each voxel of stritaum_[loss vs. no-incentive]

Raw data 2_Activation in each voxel of stritaum_[gain vs. no-incentive]

Source Code_Heritability Brain Mapping 1

Source Code_Heritability Brain Mapping 2

Source Code_Multivariate Model fitting

## References

Aarts, E, Roelofs, A, Franke, B, Rijpkema, M, Fernandez, G, Helmich, RC, & Cools, R. 2010. Striatal dopamine mediates the interface between motivational and cognitive control in humans: evidence from genetic imaging. Neuropsychopharmacology, 35(9), 1943–1951. DOI:10.1038/npp.2010.68

Baldo, BA, & Kelley, AE. 2007. Discrete neurochemical coding of distinguishable motivational processes: insights from nucleus accumbens control of feeding. Psychopharmacology (Berl), 191(3), 439–459. DOI:10.1007/s00213-007-0741-z

Beck, A, Schlagenhauf, F, Wustenberg, T, Hein, J, Kienast, T, Kahnt, T, Schmack, K, Hagele, C, Knutson, B, Heinz, A, & Wrase, J. 2009. Ventral Striatal Activation During Reward Anticipation Correlates with Impulsivity in Alcoholics. Biological Psychiatry, 66(8), 734–742. DOI:10.1016/j.biopsych.2009.04.035

Bergen, SE, Gardner, CO, & Kendler, KS. 2007. Age-related changes in heritability of behavioral phenotypes over adolescence and young adulthood: a meta-analysis. Twin Res Hum Genet, 10(3), 423–433. DOI:10.1375/twin.10.3.423

Berridge, KC. 2003. Pleasures of the brain. Brain and Cognition, 52(1), 106–128. DOI:10.1016/s0278-2626(03)00014-9

Berridge, KC. 2007. The debate over dopamine’s role in reward: the case for incentive salience. Psychopharmacology (Berl), 191(3), 391–431. DOI:10.1007/s00213-006-0578-x

Berridge, KC. 2012. From prediction error to incentive salience: mesolimbic computation of reward motivation. European Journal of Neuroscience, 35(7), 1124–1143. DOI:10.1111/j.1460-9568.2012.07990.x

Berridge, KC, & Robinson, TE. 1998. Brain Research Reviews. Brain Res Brain Res Rev, 28(3), 309–369.

Bossong, MG, & Kahn, RS. 2016. The Salience of Reward. JAMA Psychiatry, 73(8), 777–778. DOI:10.1001/jamapsychiatry.2016.1134

Braff, DL, Freedman, R, Schork, NJ, & Gottesman, II. 2007. Deconstructing schizophrenia: An overview of the use of endophenotypes in order to understand a complex disorder. Schizophrenia Bulletin, 33(1), 21–32. DOI:10.1093/schbul/sbl049

Chan, RC, Gooding, DC, Shi, HS, Geng, FL, Xie, DJ, Yang, ZY, Liu, WH, Wang, Y, Yan, C, Shi, C, Lui, SS, & Cheung, EF. 2016. Evidence of structural invariance across three groups of Meehlian schizotypes. NPJ Schizophrenia, 2, 16016. DOI:10.1038/npjschz.2016.16

Chan, RC, Li, Z, Li, K, Zeng, YW, Xie, WZ, Yan, C, Cheung, EF, & Jin, Z. 2016. Distinct processing of social and monetary rewards in late adolescents with trait anhedonia. Neuropsychology, 30(3), 274–280. DOI:10.1037/neu0000233

Chan, RC, Wang, Y, Huang, J, Shi, Y, Wang, Y, Hong, X, Ma, Z, Li, Z, Lai, MK, & Kring, AM. 2010. Anticipatory and consummatory components of the experience of pleasure in schizophrenia: cross-cultural validation and extension. Psychiatry Research, 175(1–2), 181–183. DOI:10.1016/j.psychres.2009.01.020

Chan, RCK, Shi, YF, Lai, MK, Wang, YN, Wang, Y, & Kring, AM. 2012. The Temporal Experience of Pleasure Scale (TEPS): Exploration and Confirmation of Factor Structure in a Healthy Chinese Sample. Plos One, 7(4). DOI:10.1371/journal.pone.0035352

Chan, RCK, Wang, Y, Yan, C, Zhao, Q, McGrath, J, Hsi, XL, & Stone, WS. 2012. A Study of Trait Anhedonia in Non-Clinical Chinese Samples: Evidence from the Chapman Scales for Physical and Social Anhedonia. Plos One, 7(4). DOI:ARTNe3427510.1371/journal.pone.0034275

Chen, J, Li, X, Chen, Z, Yang, X, Zhang, J, Duan, Q, & Ge, X. 2010. Optimization of zygosity determination by questionnaire and DNA genotyping in Chinese adolescent twins. Twin Research and Human Genetics, 13(2), 194–200. DOI:10.1375/twin.13.2.194

Chen, J, Li, X, Zhang, J, Natsuaki, MN, Leve, LD, Harold, GT, Chen, Z, Yang, X, Guo, F, Zhang, J, & Ge, X. 2013. The Beijing Twin Study (BeTwiSt): a longitudinal study of child and adolescent development. Twin Research and Human Genetics, 16(1), 91–97. DOI:10.1017/thg.2012.115

Craig, W. 1917. Appetites and Aversions as Constituents of Instincts. Proc Natl Acad Sci U S A, 3(12), 685–688.

Cuthbert, BN. 2014. The RDoC framework: facilitating transition from ICD/DSM to dimensional approaches that integrate neuroscience and psychopathology. World Psychiatry, 13(1), 28–35. DOI:10.1002/wps.20087

Cuthbert, BN, & Insel, TR. 2010. Toward New Approaches to Psychotic Disorders: The NIMH Research Domain Criteria Project. Schizophrenia Bulletin, 36(6), 1061–1062. DOI:10.1093/schbul/sbq108

de Leeuw, M, Kahn, RS, & Vink, M. 2015. Fronto-striatal dysfunction during reward processing in unaffected siblings of schizophrenia patients. Schizophrenia Bulletin, 41(1), 94–103. DOI:10.1093/schbul/sbu153

Docherty, AR, Sponheim, SR, & Kerns, JG. 2015. Self-reported affective traits and current affective experiences of biological relatives of people with schizophrenia. Schizophrenia Research, 161(2–3), 340–344. DOI:10.1016/j.schres.2014.11.013

Dreher, JC, Kohn, P, Kolachana, B, Weinberger, DR, & Berman, KF. 2009. Variation in dopamine genes influences responsivity of the human reward system. Proc Natl Acad Sci U S A, 106(2), 617–622. DOI:10.1073/pnas.0805517106

Forbes, EE, Brown, SM, Kimak, M, Ferrell, RE, Manuck, SB, & Hariri, AR. 2009. Genetic variation in components of dopamine neurotransmission impacts ventral striatal reactivity associated with impulsivity. Molecular Psychiatry, 14(1), 60–70. DOI:10.1038/sj.mp.4002086

Gottesman, II, & Gould, TD. 2003. The endophenotype concept in psychiatry: etymology and strategic intentions. The American Journal of Psychiatry, 160(4), 636–645. DOI:10.1176/appi.ajp.160.4.636

Grimm, O, Heinz, A, Walter, H, Kirsch, P, Erk, S, Haddad, L, Plichta, MM, Romanczuk-Seiferth, N, Pohland, L, Mohnke, S, Muhleisen, TW, Mattheisen, M, Witt, SH, Schafer, A, Cichon, S, Nothen, M, Rietschel, M, Tost, H, & Meyer-Lindenberg, A. 2014. Striatal response to reward anticipation: evidence for a systems-level intermediate phenotype for schizophrenia. JAMA Psychiatry, 71(5), 531–539. DOI:10.1001/jamapsychiatry.2014.9

Haber, SN, & Knutson, B. 2010. The Reward Circuit: Linking Primate Anatomy and Human Imaging. Neuropsychopharmacology, 35(1), 4–26. DOI:10.1038/npp.2009.129

Hahn, T, Heinzel, S, Dresler, T, Plichta, MM, Renner, TJ, Markulin, F, Jakob, PM, Lesch, KP, & Fallgatter, AJ. 2011. Association between reward-related activation in the ventral striatum and trait reward sensitivity is moderated by dopamine transporter genotype. Human Brain Mapping, 32(10), 1557–1565. DOI:10.1002/hbm.21127

Hay, DA, Martin, NG, Foley, D, Treloar, SA, Kirk, KM, & Heath, AC. 2001. Phenotypic and genetic analyses of a short measure of psychosis-proneness in a largescale Australian twin study. Twin Research, 4(1), 30–40.

Iversen, LL. 2010. Dopamine handbook. Oxford; New York: Oxford University Press.

Jenkinson, M, Beckmann, CF, Behrens, TE, Woolrich, MW, & Smith, SM. 2012. Fsl. Neuroimage, 62(2), 782–790. DOI:10.1016/j.neuroimage.2011.09.015

Kendler, KS, & Hewitt, JH. 1992. The structure of self-report schizotypy in twins. Journal of Personality Disorders, 6(1), 1–17.

Kendler, KS, Thacker, L, & Walsh, D. 1996. Self-report measures of schizotypy as indices of familial vulnerability to schizophrenia. Schizophrenia Bulletin, 22(3), 511–520.

Knutson, B, Bhanji, JP, Cooney, RE, Atlas, LY, & Gotlib, IH. 2008. Neural responses to monetary incentives in major depression. Biological Psychiatry, 63(7), 686–692. DOI:10.1016/j.biopsych.2007.07.023

Knutson, B, Fong, GW, Adams, CM, Varner, JL, & Hommer, D. 2001. Dissociation of reward anticipation and outcome with event-related fMRI. Neuroreport, 12(17), 3683–3687. DOI:10.1097/00001756-200112040-00016

Knutson, B, & Gibbs, SEB. 2007. Linking nucleus accumbens dopamine and blood oxygenation. Psychopharmacology, 191(3), 813–822. DOI:10.1007/s00213-006-0686-7

Knutson, B, & Greer, SM. 2008. Anticipatory affect: neural correlates and consequences for choice. Philosophical Transactions of the Royal Society B-Biological Sciences, 363(1511), 3771–3786. DOI:10.1098/rstb.2008.0155

Knutson, B, Westdorp, A, Kaiser, E, & Hommer, D. 2000. FMRI visualization of brain activity during a monetary incentive delay task. Neuroimage, 12(1), 20–27. DOI:10.1006/nimg.2000.0593

Kring, AM, & Barch, DM. 2014. The motivation and pleasure dimension of negative symptoms: neural substrates and behavioral outputs. Eur Neuropsychopharmacol, 24(5), 725–736. DOI:10.1016/j.euroneuro.2013.06.007

Lancaster, TM, Linden, DE, Tansey, KE, Banaschewski, T, Bokde, AL, Bromberg, U, Buchel, C, Cattrell, A, Conrod, PJ, Flor, H, Frouin, V, Gallinat, J, Garavan, H, Gowland, P, Heinz, A, Ittermann, B, Martinot, JL, Paillere Martinot, ML, Artiges, E, Lemaitre, H, Nees, F, Orfanos, DP, Paus, T, Poustka, L, Smolka, MN, Vetter, NC, Jurk, S, Mennigen, E, Walter, H, Whelan, R, Schumann, G, & Consortium, I. 2016. Polygenic Risk of Psychosis and Ventral Striatal Activation During Reward Processing in Healthy Adolescents. JAMA Psychiatry, 73(8), 852–861. DOI:10.1001/jamapsychiatry.2016.1135

Lenroot, RK, Schmitt, JE, Ordaz, SJ, Wallace, GL, Neale, MC, Lerch, JP, Kendler, KS, Evans, AC, & Giedd, JN. 2009. Differences in genetic and environmental influences on the human cerebral cortex associated with development during childhood and adolescence. Hum Brain Mapp, 30(1), 163–174. DOI:10.1002/hbm.20494

Li, Z, Huang, J, Xu, T, Wang, Y, Li, K, Zeng, YW, Lui, SSY, Cheung, EFC, Jin, Z, Dazzan, P, Glahn, DC, & Chan, RCK. 2018. Neural mechanism and heritability of complex motor sequence and audiovisual integration: A healthy twin study. Human Brain Mapping, 39(3), 1438–1448. DOI:10.1002/hbm.23935

Li, Z, Lui, SS, Geng, FL, Li, Y, Li, WX, Wang, CY, Tan, SP, Cheung, EF, Kring, AM, & Chan, RC. 2015. Experiential pleasure deficits in different stages of schizophrenia. Schizophrenia Research, 166(1–3), 98–103. DOI:10.1016/j.schres.2015.05.041

Linney, YM, Murray, RM, Peters, ER, MacDonald, AM, Rijsdijk, F, & Sham, PC. 2003. A quantitative genetic analysis of schizotypal personality traits. Psychological Medicine, 33(5), 803–816. DOI:10.1017/s0033291703007906

MacDonald, AW, Pogue-Geile, MF, Debski, TT, & Manuck, S. 2001. Genetic and environmental influences on schizotypy: A community based twin study. Schizophrenia Bulletin, 27(1), 47–58.

Meehl, PE. 1975. Hedonic Capacity - Some Conjectures. Bulletin of the Menninger Clinic, 39(4), 295–307.

Meehl, PE. 1990. Toward an Integrated Theory of Schizotaxia, Schizotypy, and Schizophrenia. Journal of Personality Disorders, 4(1), 1–99. DOI:10.1521/pedi.1990.4.1.1

Neale, MC, Hunter, MD, Pritikin, JN, Zahery, M, Brick, TR, Kirkpatrick, RM, Estabrook, R, Bates, TC, Maes, HH, & Boker, SM. 2016. OpenMx 2.0: Extended Structural Equation and Statistical Modeling. Psychometrika, 81(2), 535–549. DOI:10.1007/s11336-014-9435-8

Neale, MC, & Maes, HHM. 2004. Methodology for Genetic Studies of Twins and Families. Dordrecht, The Netherlands: Kluwer Academic Publishers.

Nikolova, YS, Ferrell, RE, Manuck, SB, & Hariri, AR. 2011. Multilocus genetic profile for dopamine signaling predicts ventral striatum reactivity. Neuropsychopharmacology, 36(9), 1940–1947. DOI:10.1038/npp.2011.82

Oswald, LM, Wong, DF, Zhou, Y, Kumar, A, Brasic, J, Alexander, M, Ye, W, Kuwabara, H, Hilton, J, & Wand, GS. 2007. Impulsivity and chronic stress are associated with amphetamine-induced striatal dopamine release. Neuroimage, 36(1), 153–166. DOI:10.1016/j.neuroimage.2007.01.055

Pizzagalli, DA. 2014. Depression, stress, and anhedonia: toward a synthesis and integrated model. Annual Review of Clinical Psychology, 10, 393–423. DOI:10.1146/annurev-clinpsy-050212-185606

Pizzagalli, DA, Holmes, AJ, Dillon, DG, Goetz, EL, Birk, JL, Bogdan, R, Dougherty, DD, Iosifescu, DV, Rauch, SL, & Fava, M. 2009. Reduced Caudate and Nucleus Accumbens Response to Rewards in Unmedicated Individuals With Major Depressive Disorder. American Journal of Psychiatry, 166(6), 702–710. DOI:10.1176/appi.ajp.2008.08081201

Power, JD, Barnes, KA, Snyder, AZ, Schlaggar, BL, & Petersen, SE. 2012. Spurious but systematic correlations in functional connectivity MRI networks arise from subject motion. Neuroimage, 59(3), 2142–2154. DOI:10.1016/j.neuroimage.2011.10.018

Radua, J, Schmidt, A, Borgwardt, S, Heinz, A, Schlagenhauf, F, McGuire, P, & Fusar-Poli, P. 2015. Ventral Striatal Activation During Reward Processing in Psychosis: A Neurofunctional Meta-Analysis. JAMA Psychiatry, 72(12), 1243–1251. DOI:10.1001/jamapsychiatry.2015.2196

Silverman, MH, Krueger, RF, Iacono, WG, Malone, SM, Hunt, RH, & Thomas, KM. 2014. Quantifying familial influences on brain activation during the monetary incentive delay task: an adolescent monozygotic twin study. Biological Psychology, 103, 7–14. DOI:10.1016/j.biopsycho.2014.07.016

Smoski, MJ, Rittenberg, A, & Dichter, GS. 2011. Major depressive disorder is characterized by greater reward network activation to monetary than pleasant image rewards. Psychiatry Research, 194(3), 263–270. DOI:10.1016/j.pscychresns.2011.06.012

Stokes, PR, Shotbolt, P, Mehta, MA, Turkheimer, E, Benecke, A, Copeland, C, Turkheimer, FE, Lingford-Hughes, AR, & Howes, OD. 2013. Nature or nurture? Determining the heritability of human striatal dopamine function: an [18F]-DOPA PET study. Neuropsychopharmacology, 38(3), 485–491. DOI:10.1038/npp.2012.207

Toulopoulou, T, van Haren, N, Zhang, X, Sham, PC, Cherny, SS, Campbell, DD, Picchioni, M, Murray, R, Boomsma, DI, Pol, HH, Brouwer, R, Schnack, H, Fananas, L, Sauer, H, Nenadic, I, Weisbrod, M, Cannon, TD, & Kahn, RS. 2015. Reciprocal causation models of cognitive vs volumetric cerebral intermediate phenotypes for schizophrenia in a pan-European twin cohort. Molecular Psychiatry, 20(11), 1482. DOI:10.1038/mp.2015.117

Tziortzi, AC, Searle, GE, Tzimopoulou, S, Salinas, C, Beaver, JD, Jenkinson, M, Laruelle, M, Rabiner, EA, & Gunn, RN. 2011. Imaging dopamine receptors in humans with [11C]-(+)-PHNO: dissection of D3 signal and anatomy. Neuroimage, 54(1), 264–277. DOI:10.1016/j.neuroimage.2010.06.044

Vignapiano, A, Mucci, A, Ford, J, Montefusco, V, Plescia, GM, Bucci, P, & Galderisi, S. 2016. Reward anticipation and trait anhedonia: An electrophysiological R. investigation in subjects with schizophrenia. Clinical Neurophysiology, 127(4), 2149–2160. DOI:10.1016/j.clinph.2016.01.006

Vink, M, de Leeuw, M, Pouwels, R, van den Munkhof, HE, Kahn, RS, & Hillegers, M. 2016. Diminishing striatal activation across adolescent development during reward anticipation in offspring of schizophrenia patients. Schizophrenia Research, 170(1), 73–79. DOI:10.1016/j.schres.2015.11.018

Wacker, J, Dillon, DG, & Pizzagalli, DA. 2009. The role of the nucleus accumbens and rostral anterior cingulate cortex in anhedonia: integration of resting EEG, fMRI, and volumetric techniques. Neuroimage, 46(1), 327–337. DOI:10.1016/j.neuroimage.2009.01.058

Wrase, J, Schlagenhauf, F, Kienast, T, Wustenberg, T, Bermpohl, F, Kahnt, T, Beek, A, Strohle, A, Juckel, G, Knutson, B, & Heinz, A. 2007. Dysfunction of reward processing correlates with alcohol craving in detoxified alcoholics. Neuroimage, 35(2), 787–794. DOI:10.1016/j.neuroimage.2006.11.043

Yacubian, J, Sommer, T, Schroeder, K, Glascher, J, Kalisch, R, Leuenberger, B, Braus, DF, & Buchel, C. 2007. Gene-gene interaction associated with neural reward sensitivity. Proc Natl Acad Sci U S A, 104(19), 8125–8130. DOI:10.1073/pnas.0702029104

